# Generalization via superposition: Combined effects of mixed reference frame representations for explicit and implicit learning in a visuomotor adaptation task

**DOI:** 10.1101/415869

**Authors:** Eugene Poh, Jordan A. Taylor

## Abstract

Studies on generalization of learned visuomotor perturbations has generally focused on whether learning is coded in extrinsic or intrinsic reference frames. This dichotomy, however, is challenged by recent findings showing that learning is represented in a mixed reference frame. Overlooked in this framework is how learning is the result of multiple processes, such as explicit re-aiming and implicit motor adaptation. Therefore the proposed mixed representation may simply reflect the superposition of explicit and implicit generalization functions, each represented in different reference frames. Here, we characterized the individual generalization functions of explicit and implicit learning in relative isolation to determine if their combination could predict the overall generalization function when both processes are in operation. We modified the form of feedback in a visuomotor rotation task to isolate explicit and implicit learning, and tested generalization across different limb postures to dissociate the extrinsic and intrinsic representations. We found that explicit generalization occurred predominantly in an extrinsic reference frame but the amplitude was reduced with postural changes, whereas implicit generalization was phase-shifted according to a mixed reference frame representation and amplitude was maintained. A linear combination of individual explicit and implicit generalization functions accounted for nearly 85% of the variance associated with the generalization function in a typical visuomotor rotation task, where both processes are in operation. This suggests that each form of learning results from a mixed representation with distinct extrinsic and intrinsic contributions, and the combination of these features shape the generalization pattern observed at novel limb postures.

**New and noteworthy:** Generalization following learning in visuomotor adaptation tasks can reflect how the brain represents what it learns. In this study, we isolated explicit and implicit forms of learning, and showed that they are derived from a mixed reference frame representation with distinct extrinsic and intrinsic contributions. Furthermore, we showed that the overall generalization pattern at novel workspaces is due to the superposition of independent generalization effects developed by explicit and implicit learning processes.

## Introduction

The details of how the motor system generalizes has been an issue of considerable interest in sensorimotor control because it can provide theoretical insights into the computational principles underlying motor learning (Poggio and Bizzi 2004; Shadmehr 2004). Specifically, the frame of reference according to which learning generalizes is critical for elucidating the internal representation of newly learned motor behavior.

Much has been learned about the internal representations of learning through adaptation tasks in which visual feedback of movement trajectories is distorted, such as that induced by a virtual device that rotates the normal relationship between the visually perceived and actual hand position during reaching movements (Krakauer et al. 2000; Pine et al. 1996). In this approach, the reference frame associated with the representation is characterized by training participants to reach with the visuomotor perturbation in a single movement direction and examining generalization to different spatial positions and limb postures. The similarity with which generalization to the untrained limb posture is linked with the environment (i.e. extrinsic reference frame) or the state of the arm (i.e., intrinsic reference frame) defines the reference frame representation associated with the learning, since these two reference frames are necessary to span interaction between the body and physical world (Buneo and Andersen 2012; McGuire and Sabes 2009; Sabes 2011; Sober and Sabes 2005).

Despite over two decades of studies employing this framework, there remains substantial disagreement regarding the characteristics of the internal representation. In visuomotor rotation tasks, Wang and Sainburg (2005) and Krakauer et al. (2000) have provided support for a representation based on movement or positions extrinsic to the end effector. They found that the accuracy of reaching movements to the learned target direction from the novel posture was maintained, despite requiring different joint rotations to attain the target. In contrast, others have reported patterns of generalization that is consistent with an intrinsic joint-based representation (de Rugy et al. 2009; Krakauer et al. 2006; Rotella et al. 2014). In an isometric visuomotor rotation task, Rotella et al. (2015) found that generalization was more complete to a target that was similar in terms of the learned joint torques and muscle activations, rather than to a target that was consistent with the extrinsic movement direction at the new limb posture.

To complicate matters, Brayanov et al. (2012) showed that, when generalization was probed across a wide distribution of movement directions at various limb postures, generalization was in fact most prominent in movement directions that were intermediate between the training direction defined according to extrinsic and intrinsic reference frames. This result was interpreted to indicate that learning is encoded in a gain-field combination of extrinsic and intrinsic movement representations. The authors further proposed that this gain-field combination can explain the apparently conflicting generalization patterns in previous studies.

At present, a mechanistic explanation for the apparent mixed reference frame underlying learning in visuomotor adaptation tasks is lacking. One possibility is that learning is not the result of a single process. Indeed, it has become increasingly clear that learning in visuomotor adaptation tasks is the result of multiple processes (Diedrichsen et al. 2010; Heuer and Hegele 2008; 2011; Huang et al. 2011; Izawa and Shadmehr 2011; Mazzoni and Krakauer 2006; Taylor et al. 2014). We have recently demonstrated that, by contrasting reports of subjects’ explicit re-aiming direction with actual reaching directions, the adaptive response during a visuomotor rotation task is made up of at least two quantifiable components: Explicit re-aiming strategies that arise from awareness of the perturbation when large task performance errors are observed and implicit motor adaptation process that is driven by sensory prediction error signals that reflect the difference between the intended and actual movement outcome (Bond and Taylor 2015; McDougle et al. 2015; 2017; Poh et al. 2016; Taylor et al. 2014). Thus, given that the overall adaptive response during a visuomotor rotation task consists of both explicit and implicit forms of learning, then the ensuing generalization should carry signatures of both forms of learning, which may be very different.

The dual operation of explicit and implicit learning in a typical visuomotor rotation task makes for interesting interactions between both forms of learning, which may ultimately influence the pattern of generalization across different limb postures. Recent studies have suggested that the manner in which implicit learning generalizes to other movement directions, within a workspace, may be intimately linked to the locus of the explicit aiming direction (Day et al. 2016; McDougle et al. 2017). For example, McDougle et al. (2017) found that implicit learning was expressed maximally at the subject’s explicit aiming location, rather than the actual trained target location. If the locus of the explicit aim was maintained in extrinsic space across different limb postures, then appropriate generalization should be observed at the location of aim that is consistent in an extrinsic reference frame at the untrained limb posture. This would give the appearance of a generalization that is intermediate between extrinsic and intrinsic reference frames, as observed by Brayanov et al. (2012). This raises a more fundamental issue, that is whether the generalization in the proposed mixed reference frame when examined across different limb postures is in fact a superposition of generalization effects driven by independent effects from both explicit and implicit forms of learning, each of which may operate in separate reference frames.

Multiple lines of evidence suggest that explicit and implicit forms of learning have different patterns of generalization, and accordingly different representations. For example, Heuer and Hegele (2008; 2011) showed that, when subjects in a standard visuomotor rotation task are prompted that the perturbation is removed, aftereffects, a measure of implicit learning, generalize narrowly within the vicinity of the trained target. In contrast, explicit learning, which is quantified by a subject’s verbal reports of the direction of movement required to compensate for the perturbation generalize broadly to movements in all directions (McDougle et al. 2017; Poh et al. 2016; Heuer and Hegele 2011). Importantly, when generalization is probed across different limb postures, explicit re-aiming displayed complete transfer to the novel limb posture, whereas implicit aftereffects was significantly reduced or hardly present (Heuer and Hegele 2011). This result shows that both forms of generalization have different attributes, and thus in theory could be represented in different reference frames. However, given the broad generalization of explicit learning and the absence of implicit aftereffects at novel limb postures, the effects of generalization according to the extrinsic or intrinsic reference frame are indistinguishable. This presents a key obstacle in determining which reference frames contribute to the representation of explicit and implicit learning, and hence the question of which reference frame underlies both forms of learning has yet to be systematically addressed.

Here we designed a series of visuomotor rotation experiments to characterize and quantify the effects of explicit and implicit learning in relative isolation, and then examined generalization across different limb postures to dissociate the extrinsic and intrinsic reference frames of representation. To selectively engage explicit learning, subjects performed a visuomotor rotation task with delayed endpoint feedback, which has been shown to severely disrupt development of implicit learning, but not explicit re-aiming strategies in a visuomotor rotation task (Brudner et al. 2016; Schween and Hegele 2017). This presents a particularly promising approach, in lieu of using verbal re-aiming reports as a proxy for explicit learning, because it allows re-aiming strategies to develop naturally in the task without visual cues or verbal instruction. What’s more, it may also allow for an unbiased estimate of the generalization function, as previous attempts to measure the generalization function have relied on visual cues or post-tests that may warp the generalization function itself (Heuer and Hegele, 2008, 2011; McDougle et al 2017).

To selectively engage implicit learning, we introduced a task-irrelevant-visual-error clamp wherein the cursor followed an invariant trajectory regardless of the subject’s movement direction. Critically, the clamped visual feedback introduced a constant visuomotor discrepancy between angular hand position and visual feedback throughout adaptation, which rendered the visual error signals observed irrelevant to the task. By emphasizing that the task goal is to reach for the target irrespective of the direction of the clamped visual feedback, this approach is thought to suppress explicit re-aiming strategies, and any changes in behavior are presumed to reflect the signature of an implicit form of learning (Morehead et al. 2017).

The goal here was to isolate each learning process, characterize the generalization function across different limb postures, and then determine if the combination of the independent generalization functions could predict the overall generalization function when both processes were in operation in a standard visuomotor rotation task. We hypothesized that explicit and implicit learning would display generalization in different reference frames, and their superposition could account for the discrepancies between prior studies and the apparent mixture of reference frames representations. Contrary to our initial hypothesis, we found that both explicit and implicit learning results from mixed representations, but each form of learning had differing relative mixtures of extrinsic and intrinsic representation.

## Methods

### Subjects

A total of 48 subjects (16 males, 32 females; age range: 18-32 years old) were recruited from the research participation pool of the Department of Psychology at Princeton University in exchange for course credit or money. The sample size was determined by a power analysis (alpha = 0.95), which sought to replicate the effect size of a relevant result (*d=1.6*; shift of generalization pattern from workspace 1 to workspace 2 in Experiment 1 in Brayanov et al. 2012). The required sample size was 15, but to counterbalance training target directions and rotation signs, we recruited a sample of 16 subjects for each condition. All subjects were right-handed as verified with the Edinburgh handedness inventory (Oldfield 1971) and provided written informed consent before participation. The protocol was approved by Princeton University’s Institutional Review Board.

### Apparatus and task

Subjects sat comfortably on a chair and made planar reaching movements while holding onto the handle of a 2-link robotic manipulandum (KINARM End-Point, BKIN Technologies). The standard configuration of robot’s handle was inverted and the subject’s right arm was supported against gravity using a ceiling-mounted sling such that arm movements occurred predominantly in the horizontal plane. All visual stimuli were projected from a 47-inch 1920 by 1080-pixel resolution display (LG47LD452C, LG Electronics), which was horizontally mounted and inverted to face a mirror that was 6cm above the robot’s handle. The mirror occluded the manipulandum, and the hand and arm of the subject, preventing direct visual feedback of the hand location. 2-D position of the hand location was sampled at 1000Hz.

In all experiments, subjects performed 7.5cm point-to-point reaching movements from two limb postures at the left, training workspace (W1) and the right, transfer workspace (W2; Figure 1A). These workspaces were uniquely defined for each subject based on their measured forearm and upper arm lengths. In W1, a subject’s shoulder and elbow angles were 75° and 90° respectively. In W2, the subject’s elbow angle was maintained but the shoulder angle was rotated clockwise 45°. Note, a greater angular separation between W1 and W2 allows for better dissociation between extrinsic and intrinsic reference frames; however, angles greater than 45° greatly affect the viewing angle and joint mobility. The reaching targets (24 target positions regularly spaced 15° around a circle) were located on the circumference of an invisible circle (radius: 7.5cm) centered on a starting location defined by W1 and W2.

**Figure 1.**
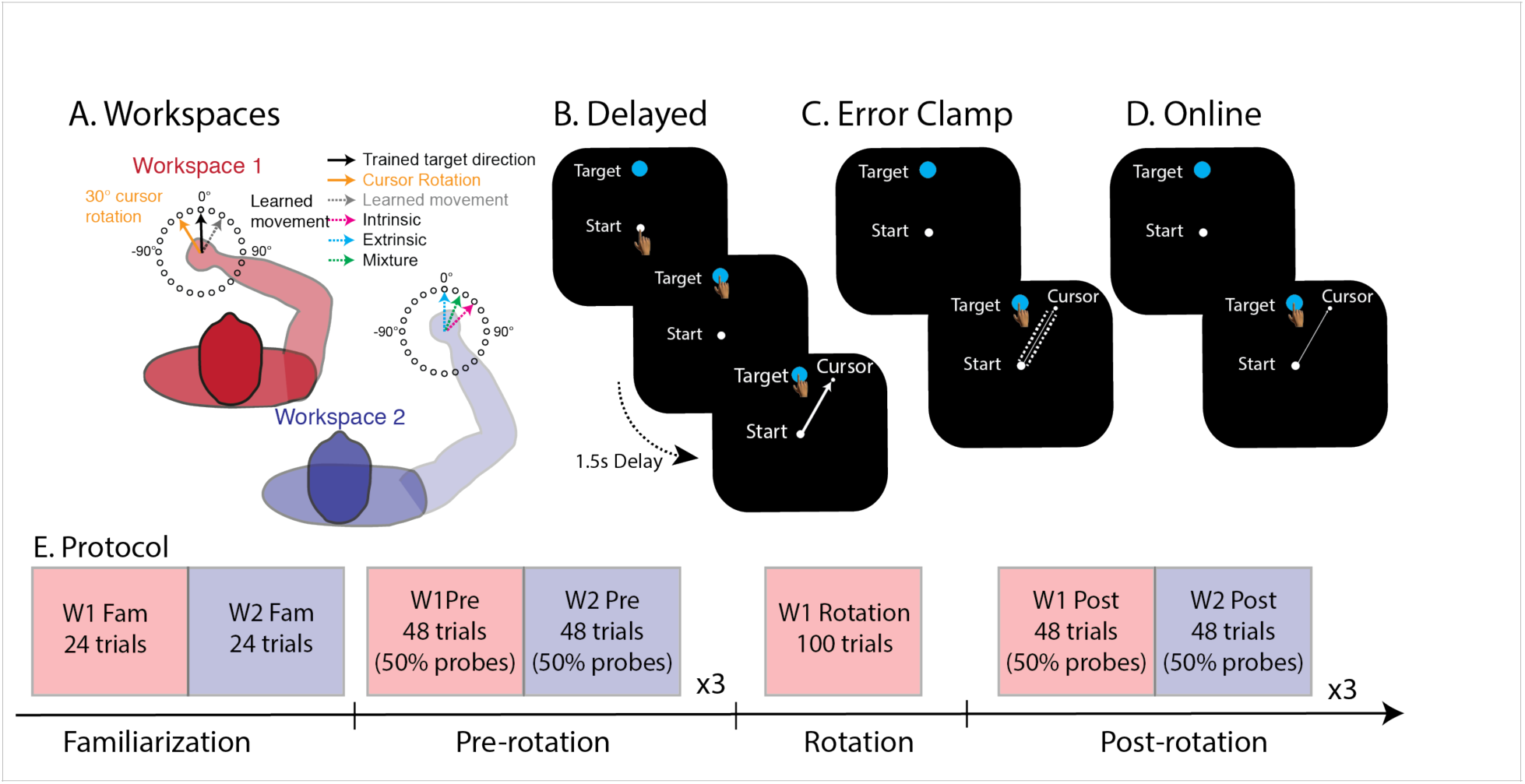
Experimental set up and conditions. A. Subjects learned to compensate for a 30° rotation at workspace 1 (W1) before generalization was tested to 24 targets in workspace 2 (W2). The two workspaces were separated by a 45° clockwise rotation of the shoulder to dissociate the extrinsic and intrinsic representations. B. In the explicit condition, subjects received delayed endpoint feedback of their performance during training. C. In the implicit condition, the feedback cursor was clamped during training, such that it followed an invariant trajectory that is offset by ±30° regardless of the subject’s movement direction. D. In the combined condition, subjects learned to overcome the visuomotor rotation using online visual feedback. This approach has been shown to allow operation of both explicit and implicit forms of learning. E. Training protocol.

Each trial began when the robotic manipulandum positioned the subject’s hand within a starting location (grey empty circle: radius: 0.5cm). When the hand was within the starting location, the grey circle turned white and a white cursor (white filled circle: radius: 0.5cm) representing the current hand position appeared. The hand had to remain in the starting location for 300ms before a target (blue filled circle: radius: 0.5cm) appeared at one of 24 possible target locations (regularly spaced 15° around a circle). The presentation of each target coincided with a beep to initiate the trial and subjects were instructed to make a ballistic movement that “sliced” through the target.

Depending on the particular training condition in W1 (see below), subjects either received delayed endpoint feedback or continuous online feedback. For delayed endpoint feedback trials, visual feedback of the cursor position was eliminated once movement velocity exceeded 5cm/s. Endpoint feedback of the cursor reappeared at the terminal position of movement 1.5s after radial movement amplitude had exceeded 7.5cm. For continuous online feedback trials, feedback of cursor location remained visible for the duration of the outbound movement. When radial movement amplitude exceeded 7.5cm, the feedback cursor froze on the screen for one second. The blue target changed to green if the feedback cursor overlapped any part of the target, otherwise the target turned red. After each trial, the target and cursor were extinguished and the robotic manipulandum repositioned the limb at a starting location defined by the training or transfer limb posture to start the next trial.

### Experimental Procedures

Subjects were pseudo-randomly assigned to one of three training conditions: explicit, implicit and combined conditions. Specifically, the explicit (n=16) and implicit conditions (n=16) were aimed at characterizing the pattern of generalization for explicit and implicit forms of learning in relative isolation, which was accomplished by providing delayed endpoint feedback and clamped visual feedback, respectively, during a visuomotor rotation task. While, the goal of the combined condition (n=16) was to determine whether the generalization function that emerges in a standard visuomotor rotation task, when both implicit and explicit forms of learning are operating simultaneously, could be accounted for by the combination of separate implicit and explicit generalization functions obtained.

Each condition began with a brief familiarization phase, which consisted of a block of 24 trials (one to each target location) in W1 and W2 (Fig. 1E). During these movements, subjects in the explicit condition received delayed endpoint feedback of their movements while subjects in the implicit and combined condition received online cursor feedback. Following familiarization, in the second pre-rotation phase, subjects completed 144 trials, divided into three blocks of 48 trials, in each workspace. During each block, visual feedback of the movements was only presented on 50% of the trials. Specifically, after every two trials with delayed endpoint feedback in the explicit condition, or online cursor feedback in the implicit and combined conditions, two probe trials without any visual feedback was presented to assess baseline performance at a workspace. We alternated between W1 And W2 after each block of 48 trials (24 feedback trials, 24 probe trials for each block). This resulted in three probe trials at each target location in both W1 and W2.

In the subsequent rotation phase, a 30° visuomotor rotation was introduced to a single training movement direction (selected from eight possible locations: 0°, 45°, 90°, 135°, 180°, 225°, 270°, 315°; counterbalanced across subjects) for 100 trials at the training target in W1. Note that the size of visuomotor rotation was chosen to match the perturbation used in a prior study (Brayanov et al., 2012). The direction of the 30° rotation was also counterbalanced across subjects.

#### Explicit condition

For the explicit condition, subjects received rotated delayed endpoint feedback of their performance (Figure 1B). Critically, this approach has been shown to be effective in limiting the development of implicit learning, and appears to result in predominantly explicit forms of learning during a visuomotor rotation task (Brudner et al., 2016; Schween and Hegele 2017).

#### Implicit condition

For the implicit condition, subjects experienced clamped visual feedback (Figure 1C): The cursor followed an invariant trajectory that is offset by ±30° (n=8: clockwise; n=8: counterclockwise) regardless of the subject’s movement direction (Morehead et al. 2017). Because subjects were instructed to ignore the clamped visual feedback and to move their hand directly to the target, this paradigm should suppress any explicit re-aiming strategies since the visual error signals observed is irrelevant to the task. Thus, any changes in behavior would reflect learning from implicit motor adaptation (Morehead et al. 2017).

#### Combined condition

For the combined condition, subjects experienced rotated online visual feedback at the training target in W1 (Figure 1D). This approach allows both explicit and implicit learning to operate during training (Poh et al., 2016; McDougle et al., 2015 & 2017; Bond and Taylor 2015; Taylor et al. 2014).

Finally, to examine the pattern of generalization in the post-rotation phase, subjects performed a total of 72 probe trials with no visual feedback at each limb posture (3 probe trials for each of the 24 movement directions). After every two trials with no visual feedback, the perturbed visual feedback was restored during movements made to the training target in W1 for the next two trials to maintain learning.

### Data analysis

#### Behavioral measure

Terminal reach directions were computed as the direction of the vector connecting the subjects’ hand location at movement onset to the point when movements passed a radial distance of 7.5 cm. Movement onset was defined when the movement speed threshold first exceeded a speed of 5cm/s. For all trials, we computed the directional errors by calculating the angular difference between the direction of target and terminal reach directions. To examine changes in movements made between workspaces W1 and W2, we subtracted the directional errors for each target during probe trials in the post-rotation phase from the directional errors in the pre-rotation phase for each target to yield the change in hand angle from baseline for each target in each workspace. This was done to mitigate any potential performance biases due to biomechanical differences across workspaces and target locations. All performance measures are reported as means ±SEM unless otherwise stated.

#### Measuring generalization

To characterize the generalization pattern in W1 and W2, the changes in reach direction for each target location defined according to an extrinsic Cartesian reference frame were fit with Gaussian tuning functions (Brayanov et al. 2012; Tanaka et al. 2009):

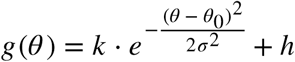

The generalization function, g(*θ*) is centered on the target direction eliciting the greatest change in reach direction (*θ*_0_), has an amplitude of *k*, and is local with a width characterized by (*σ*). The generalization function also contains a constant offset parameter, *h,* which has been thought to reflect the global (uniform) portion of motor adaptation (see Brayanov et al. 2012). The amplitude of the fit was constrained to the maximum change in reach direction at W1. We fit the gaussian function to individual subject’s data for each condition at the training and transfer limb postures to estimate the amplitude (*k*), centre (*θ*), width (*σ*) and vertical offset (*h*) of the generalization at W1 and W2. The parameters of the generalization function were estimated using the least squares method of the function “fit” in MATLAB.

#### Defining the intrinsic and extrinsic space

The purpose of this study is to examine how the extrinsic and intrinsic reference frames contribute to the internal representation of explicit and implicit forms of learning. To achieve this, we dissociated the extrinsic and intrinsic reference frames of representation by changing the limb postures relative to the training limb posture. To parameterize each movement with respect to the extrinsic and intrinsic reference frame for which we tested generalization, we adopted the framework of Brayanov et al. (2012), wherein it is assumed that a movement can be defined according to reference frames that are either extrinsic or intrinsic relative to the spatial position of the limb. Briefly, in an extrinsic Cartesian reference frame, a straight point-to-point movement can be characterized by a vector that connects the movement starting and end points. Figure 2A shows the movement vectors in Cartesian space for all 48 target positions in the training and transfer limb postures. The start locations for each of the limb posture depicted in Figure 2A were calculated based on the forearm and upper arm lengths averaged across all 48 subjects. In this conception, the relationship between any two straight movements with the same origin but ending at two different targets is defined by the angular distance between the movement vectors in Cartesian space. Correspondingly, the same two movements can be defined in an intrinsic joint-based reference frame as a pair of joint-space movement vectors that share a common origin. The joint-space angle separating the the two vectors describes the angular separation between the two movements in an intrinsic joint-based reference frame. Figure 2B shows the intrinsic joint-based representations of the movements toward each target direction at both workspaces. Here, we computed the intrinsic joint-based trajectories using the inverse kinematics equations for a two-link robotic manipulandum (Spong 2006). Shoulder and elbow angles used for calculations were defined relative to the frontal plane.

**Figure 2.**
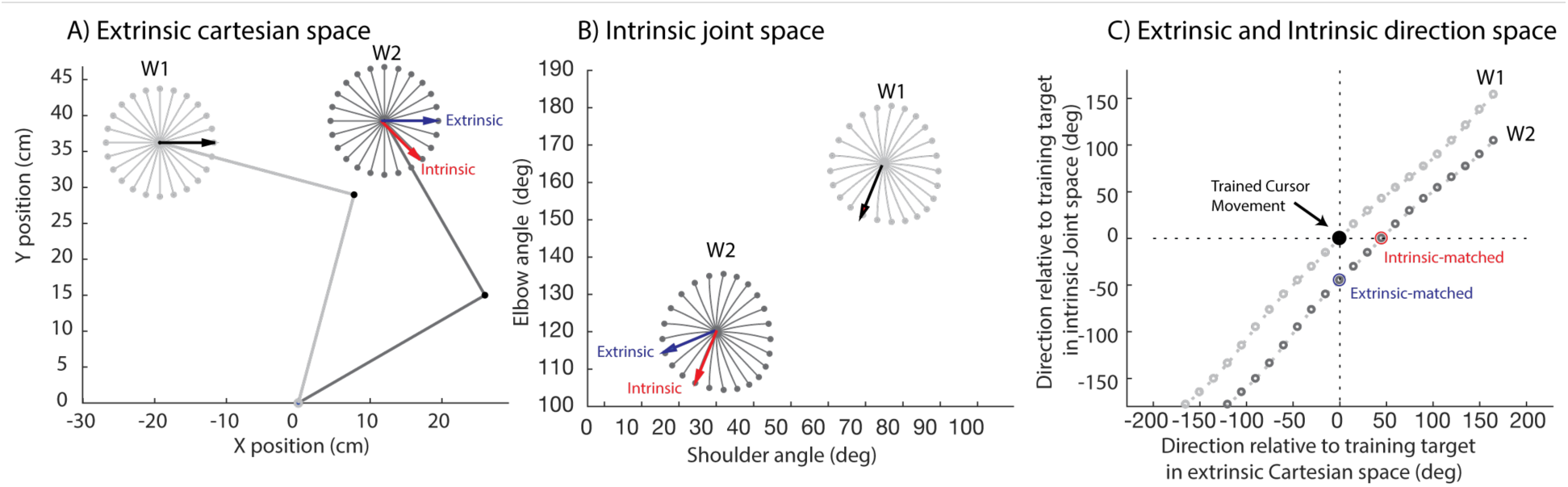
Defining the intrinsic and extrinsic movement representations. A) Extrinsic movement representations of ideal cursor movements. In this example, subjects adapted to a visuomotor rotation to the 0° target at W1 (black arrow), and generalization was probe to an array of 24 targets (regularly spaced at 15°) at both W1 and W2. The grey trajectories illustrate the ideal cursor movements to all targets in W1 and W2. In W2, the blue arrow illustrates the trained cursor movement in extrinsic Cartesian space, while the red arrow shows the trained cursor movement in intrinsic joint space. It is important to note that the parallel blue and black arrows in A indicate that these two movements require the same change in extrinsic Cartesian space. B) Intrinsic movement representations of ideal cursor movements. Here, the x-axis represents the the shoulder angle relative to the torso, while the y-axis represents the elbow angle relative to the torso. In W1, the black arrow represents the trained direction in joint space. Similarly, in W2, the blue arrow illustrates the trained cursor movement in extrinsic Cartesian space, while the red arrow shows the trained cursor movement in intrinsic joint space. In contrast to panel A, the black and red arrows are now parallel to each other, implying that those two movements require the same joint rotations. C) Distance of the targets in extrinsic and intrinsic space. On the x-axis, the distance between different targets is calculated as the difference between that and the trained target in extrinsic Cartesian space. In comparison, the y-axis depicts the distance between each target and the trained one as the difference between that and the trained target in intrinsic joint space.

To determine the angular distance between the trained movement direction and any given movement direction in the extrinsic reference frame, we computed the difference between the trained and the testing target location in terms of the change in x-y coordinates:

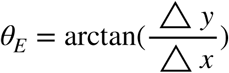

In comparison, the angular distance between the trained movement direction and any given movement in the intrinsic reference frame (*θ*_*I*_) was defined in terms of the change shoulder-elbow coordinates:

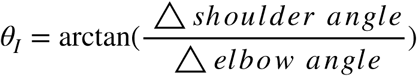

Figure 2C illustrates the two dimensional extrinsic-intrinsic space employing the two reference frames as cardinal axis in a two-dimensional plot. This plot characterizes the distance in both the extrinsic and intrinsic reference frames between the trained movement and any given movement direction for which generalization was tested. Note that the trained target in the training limb configuration is located at the origin (0°, 0°). The limb posture manipulation in our study that resulted in a 45° rotation in the shoulder angle between W1 and W2 produced a 45° shift in the locus of the target locations (Figure 2C). This shift is such that the target location corresponding to the trained movement direction in intrinsic space is located at position (-45°, 0°), whereas the target location corresponding to the trained movement direction in extrinsic space is located at (0°, 45°).

#### Generalization predictions

The manner in which the pattern of generalization is expressed at W2 depends on the reference frame according to which learning is represented. In our study, subjects learned to compensate for a visuomotor rotation in a single movement direction at W1, and generalization was tested at W2 with the shoulder angle rotated 45° clockwise. Based on the extrinsic-intrinsic space defined above, if learning mapped exclusively onto an intrinsic reference frame, the direction of peak generalization would be observed at the target direction is rotated 45° clockwise relative to the trained target direction defined extrinsically at W2 (W2-W1 ≈ 45°). In contrast, if learning is mapped exclusively onto an extrinsic reference frame, the pattern of generalization at the W1 and W2 should be identical. In this case, the direction of peak generalization should remain invariant in extrinsic space, irrespective of limb posture (W2≈W1). Additionally, we also considered the possibility that the representation involve a mixture of reference frames. Depending on the mixture ratios between extrinsic and intrinsic reference frames, this would cause the pattern of generalization to be shifted to an intermediate position between the trained target direction defined intrinsically and extrinsically (0 < W2-W1 < 45°). The difference between estimated peak direction at W1 and W2 were tested using single sample t-tests with the test variable set at zero or 45° in accordance with the pure extrinsic or pure intrinsic representation. Significance level for all tests was set at 0.05.

To further consider a mixed representation, we determined the relative contributions of extrinsic and intrinsic representations to the pattern of generalization using the multiplicative gain-field combination model of Brayanov et al. (2012). This model posits that the generalization to a given target direction depends on combined distance across the extrinsic and intrinsic space from the trained direction.

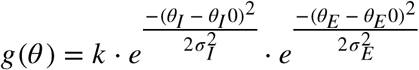

where 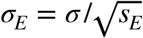 and 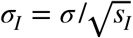. The coefficients *s*_*E*_ and *s*_*I*_ where *s*_*e*_ + *s*_*I*_ = 1, are included to allow for differential weighting of extrinsic and intrinsic components of the overall distance, and allowed us to determine the extent to which extrinsic and intrinsic reference frames contribute to the generalization at W2. Note that we used the data from W1 to constrain the width of generalization.

Because the goodness of fit to the individual generalization functions is prone to variability inherent within each subject’s data, we performed a bootstrap test with 2000 iterations to estimate the confidence intervals of the parameters estimated from the fit to the mean data from each condition. This approach allows us to determine the variability associated with the parameters of the fit to the mean data. Specifically, on each iteration, we selected 16 subjects with replacement from the respective condition and computed the generalization function and corresponding fits based on the data averaged across the selected subjects. The 95% confidence intervals were then estimated from the distribution of these fit parameters.

To statistically compare whether the distributions of the fitted parameters at W1 were different from that at W2, we used the p-value computed from the bootstrap. That is, we first calculated the difference in the estimated parameters between the training and transfer limb posture on each iteration of the bootstrap. Using the distribution of bootstrapped differences, we estimated the number of samples in the distribution that are above the absolute value of the null hypothesis. The percent of samples that are above the absolute value of the null hypothesis (0) constitute the p-value of the bootstrap. Significance level was set at 0.05.

## Results

### Learning delayed endpoint feedback and clamped visual feedback

We first set out to experimentally isolate explicit and implicit forms learning, and characterize the generalization function across different limb postures. To do this, subjects learned to compensate for a 30° visuomotor rotation with delayed endpoint or clamped visual feedback designed to isolate explicit and implicit forms of learning respectively. Both groups displayed a rapid change in hand angle in response to the visuomotor rotation at the training posture (W1; Figure 3), which is likely attributable to the fact that there is only one training target and is on par with previous studies (Bond and Taylor 2015; McDougle et al 2015, 2017). Final performance in both explicit and implicit conditions, on average, reached asymptotic levels within the 20 trials. The asymptotic performance averaged over the last 10 trials was 26.2±1.5° in the explicit condition versus 15.7±2.2° in the implicit condition. Note, while not central to the primary goal of this study — characterizing the contribution of explicit and implicit learning to the pattern of generalization — we sought to compare the asymptotic level of learning between explicit and implicit learning. We find that the delay condition resulted in significantly greater change in reach directions than the clamp condition (*p<0.001;* independent sample t-test). This difference between presumed explicit and implicit conditions are in line with previous studies which showed that explicit forms of learning are able to flexibly adjust and fully compensate for the size of rotation (Bond and Taylor 2015; Taylor et al. 2014; Brudner et al. 2016), while implicit forms of learning appears to asymptote at 15-20° (Morehead et al. 2017; Bond and Taylor 2015). This also suggests qualitatively different forms of learning when using clamped visual feedback and delayed endpoint feedback during exposure to a visuomotor rotation.

**Figure 3.**
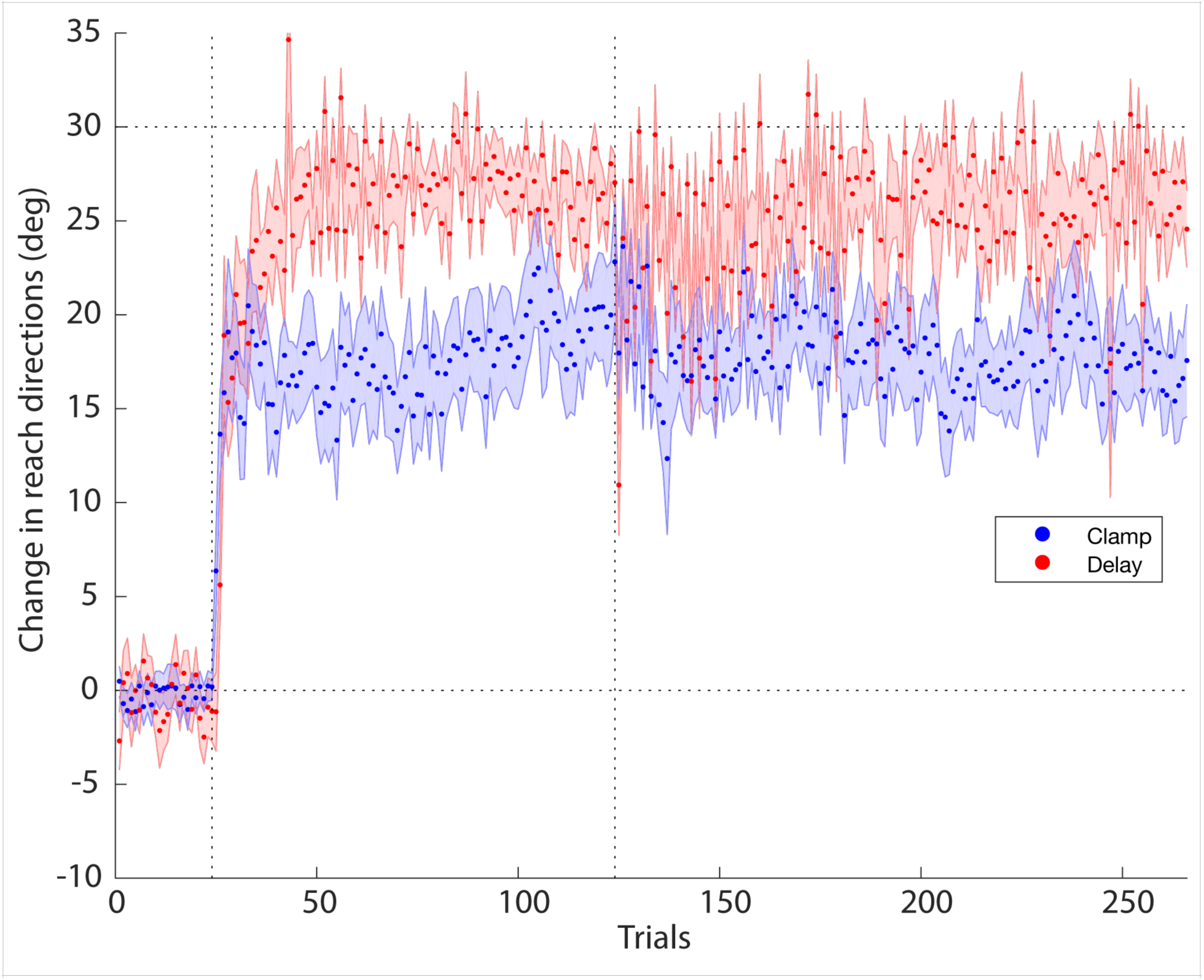
Rotation training performance for explicit and implicit conditions. Mean change in reach directions when participant trained with a cursor that was aligned (Pre-rotation phase) or rotated 30° relative to hand movement (Rotation and Post-rotation phase). The red line corresponds to reach directions in the explicit condition, while the blue line corresponds to the reach directions in the implicit condition. Error bars represent standard error of the mean.

### Generalization of explicit and implicit learning

In the subsequent post-rotation phase, probe trials without visual feedback were used to assess the pattern of generalization at W1 and W2. To quantify the change in reach direction following rotation training, the directional errors from the pre-rotation phase were subtracted from the post-rotation phase at each target location in each workspace. Subsequently, a Gaussian function with four free parameters (center position, amplitude, width and vertical offset) was fit to each individual subject’s generalization data to approximate the pattern of generalization.

Figure 4 shows the generalization of rotation training, along with the Gaussian fit, averaged across subjects in the explicit (Fig. 4A) and implicit (Fig. 4B) conditions at W1 and W2. Contrary to previous studies, which have shown quite broad generalization functions for explicit learning (Heuer and Hegele 2008; McDougle et al. 2017; 2011), here, we found it to be relatively narrow and Gaussian in shape. The explicit generalization pattern in W1 was centered near the 0° training direction (-3.4±8.8°) and has an amplitude of 22.3±2.2°, which falls short of 30° required to fully compensate for the visuomotor rotation. Further, the width of the explicit generalization pattern was 29.2±5.4° and had an offset of 4.8±2.2°.

**Figure 4.**
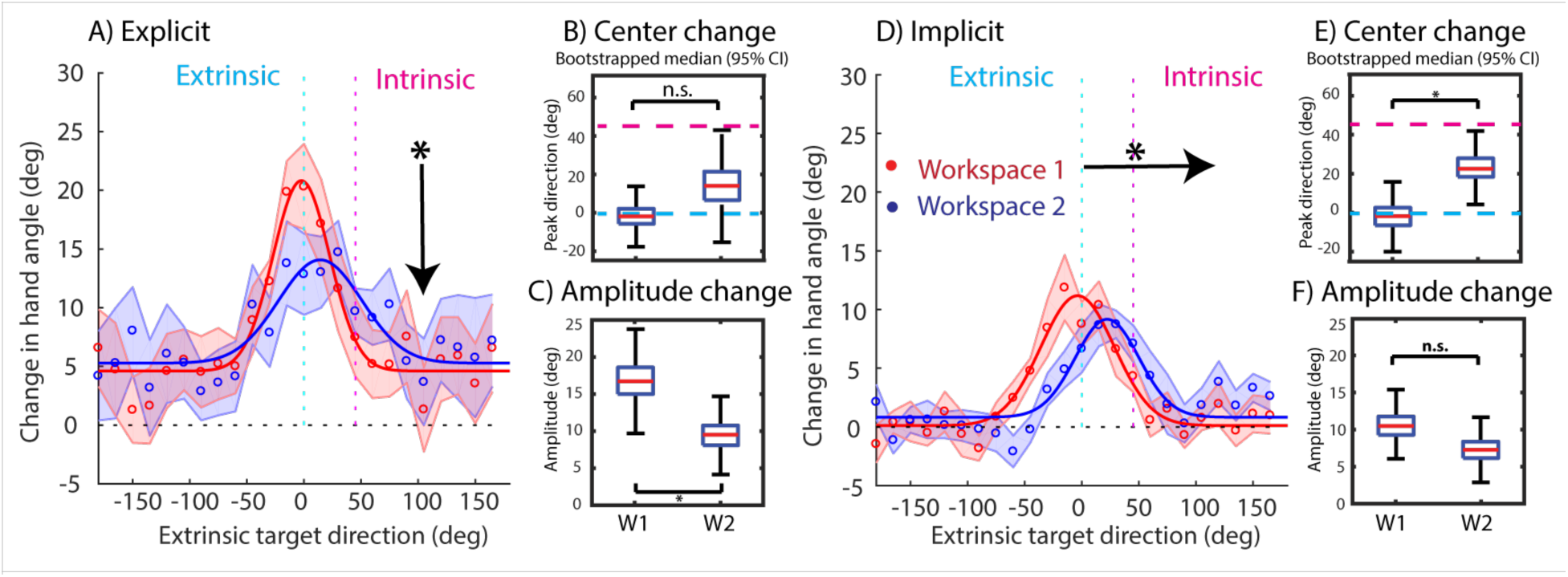
Gaussian fits for the generalization pattern in the explicit (A) and implicit (D) condition. The observed generalization pattern was the gaussian fit to the data averaged across all participants in each condition at W1 (red line) and W2 (blue line). The shaded area represents the mean±SE. B and C: Box-plots of the center of generalization (B) and amplitude (C) estimated by the gaussian fit for each of the 2000 bootstrap samples for the explicit condition at W1 and W2. E and F: Box-plots of the center of generalization (E) and amplitude (F) estimated by the gaussian fit for each of the 2000 bootstrap samples for the implicit condition at W1 and W2. Quartiles and confidence intervals indicated on the box plots. *Significant main effect following an bootstrap test at P <0.05.

Similarly, we found that the generalization function for implicit learning at W1 was relatively narrow and well-described by a Gaussian function, consistent with prior studies (McDougle et al. 2017, Morehead et al., 2017). As expected, the center location of the implicit generalization pattern in W1 was near zero (2.1±10°) with an amplitude of 13.8±1.6°, corresponding to approximately 46% compensation of the imposed visuomotor rotation. In addition, the width of the implicit generalization pattern was 25.8±4°, and there appears to be no vertical offset (0.58±0.18°).

Comparison of the generalization pattern between explicit and implicit conditions at W1 revealed that the center of the generalization pattern for the explicit (t(15) = 0.7; p=-0.4) and implicit (t(15) = 0.2; p=0.84) conditions were not significantly different from training target direction at 0° (t(15) = 0.9; p=0.39), and they were also not different from one another (t(15) = 0.32; p=0.77). Similarly, there was also no difference between the width of the generalization pattern in the explicit and implicit conditions (t(15) = 0.5; p=0.63). In contrast, the amplitude (t(15) = 3; p=0.005) was significantly smaller for the implicit condition than for the explicit condition. These results show that explicit and implicit forms of learning driven by delayed endpoint feedback and clamped visual feedback respectively, generalize locally at the trained movement direction, but that the amplitude of generalization was significantly less for the implicit condition.

Further, we note that the vertical offset in the explicit generalization function was marginally larger than the implicit generalization function (t(15) = 1.9; p=0.07). Such vertical offsets resembles the positive DC shift across the measured generalization function observed in previous visuomotor generalization studies (Krakauer et al. 2000; McDougle et al. 2017; Pearson et al. 2010; Taylor et al. 2013). Our findings imply that such DC shifts in the generalization function appear to be primarily due to explicit and not implicit learning.

We then sought to determine if the generalization functions associated with explicit and implicit learning changed when shifting to a different limb posture in W2. For the explicit condition, the Gaussian fit to each individual subject data remained centered near the extrinsic training direction at 3.3±11.3° in W2. This corresponds to a mean shift of 6.6±13.9°, which was not significantly different from the zero phase shift predicted by a pure extrinsic representation (t(15) = 0.463; p=0.325). What’s more, we found a significant drop in the amplitude (-6.4±2.7°; p=0.03), but there were no significant changes in the width (t(15) = 1.6; p=0.1) or vertical offset (t(15) = -1.2; p=0.2) of the generalization function. Taken together, these results suggest that, while the direction of peak generalization for explicit learning was maintained in an extrinsic reference frame, the amount of explicit learning acquired in W1 was significantly reduced in W2.

For the implicit condition, the Gaussian fit to individual subjects’ generalization data in W2 was centered at 26±9.7°, which represents a mean phase shift of 23.9±8.7° between W1 and W2. Interestingly, the mean phase shift is approaching half way between the zero phase shift predicted by a pure extrinsic representation (W2-W1 ≈ 0°) and the 45° shift predicted by a pure intrinsic representation (W2-W1 ≈ 45°). This phase shift in implicit generalization function at W2 is significantly different from the amount of shift predicted by the pure extrinsic or intrinsic representations (t(15) = 2.8; p=0.007 and t(15) = -2.4; p=0.014 respectively). Finally, while there was a slight but insignificant drop (-1.2±1.6°, p=0.49) in the amplitude of the generalization function, the width (t(15) = -0.04; p=0.96) and vertical offset (t(15) = -0.15; p=0.88) of the generalization function was maintained between W1 and W2. These results demonstrate that, while the direction of peak generalization for implicit learning is sensitive to limb posture, the degree of generalization is quite stable.

On average, individual subject’s gaussian fit to the pattern of explicit (R^2^= 0.37±0.04) and implicit (R^2^= 0.34±0.04) generalization was relatively poor because of the variability inherent within individual subjects’ performance. Thus, we performed a bootstrap analysis to determine the robustness of the model parameters estimated from the Gaussian fit. First, we examined the confidence intervals for the centre of generalization in W1 and W2. For the explicit condition, the median position of the generalization pattern for W1 and W2 was centered at -2° [95% confidence interval (CI) = -13.2°, 8.5°] and 14° (95% CI = -7.7°, 35°), respectively. We then calculated the p-value based on the bootstrap samples and found that they were not significantly different from one another (+15.8° bootstrap median; 95% CI = -7.7° 39.3°; p=0.09). In comparison, the median position of the generalization pattern for implicit learning in W1 and W2 was -1.8° (95% CI = -14.9°, 11.2°) and 22.7° (95% CI = 10.9°, 64.5°). Based on the bootstrap p-value, this corresponded to a significant shift of +25.2° (95% CI = 7.3° 70°; p=0.005) for the implicit generalization function.

On the other hand the median amplitude of generalization for explicit learning at W1 and W2 was 16.7° (95% CI = 11.3° 21.7°) and 9.5° (95% CI = 5.8°, 14.1°) respectively. This corresponded to a significant drop of -7.4° (95% CI = -13.6° 0.4°; p=0.03). For implicit learning, the median amplitude of generalization at W1 and W2 was 10.5° (95% CI = 7.4° 15°) and 3.8° (95% CI = 0.1°, 7.3°), which was not significantly different from each other (-3.3° bootstrap median; 95% CI = -8.7° 1.3°; p=0.08).

Next, we compared the widths and vertical offsets for the generalization pattern at W1 and W2. Consistent with the t-tests performed above, the bootstrap analysis revealed no significant changes in the widths and vertical offsets between the generalization pattern at W1 and W2 for both explicit and implicit learning. For explicit learning, the median width of the generalization function at W1 and W2 was 24.3° (95% CI = 17.5°, 34.7°) and 36.3° (95% CI = 18.2°, 74.3°) respectively (p=0.11), while the offset at W1 and W2 was 4.4° (95% CI = 0° 9.9°) and 4.8° (95% CI = 0° 11.9°), respectively (p=0.46). For implicit learning, the median width of the generalization function at W1 and W2 was 29.9° (95% CI = 22.4° 34.8°) and 27.8° (95% CI = 17.7° 66.9°) respectively (p=0.38), while the offset was 0.02° (95% CI = 0°, 1.2°) and 0.8° (95% CI = 0°, 2.1°) at W1 and W2, respectively (p=0.2).

Finally, we analyzed the goodness of fits on the bootstrap samples. For explicit learning, the median R^2^ were 0.84 (95% CI = 0.6, 0.93) and 0.62 (95% CI = 0.23, 0.85) in W1 and W2 respectively. For implicit learning, the median R^2^ were 0.83 (95% CI = 0.64, 0.91) and 0.62 (95% CI = 0.23, 0.81) in W1 and W2 respectively. Taken together, these results suggest that the phase shifts in the pattern of generalization observed for implicit learning, and the reduction in the amplitude for explicit learning, was robust, despite the presence of noise within single-subject data.

### Mixture of extrinsic and intrinsic reference frames for explicit and implicit learning

The preceding analysis on the direction of peak generalization revealed that implicit (+25.2° bootstrap median) and explicit (+15.8° bootstrap median) generalization patterns displayed different amounts of phase shifts, suggesting that there could be differential contributions of extrinsic and intrinsic reference frames. To determine the mixture ratios of extrinsic and intrinsic reference frames for generalization of explicit and implicit learning, we first fit the subject-averaged generalization pattern to the gain-field model from Brayanov and colleagues (2012), which allows for differential weighting of extrinsic and intrinsic reference frames (Fig 5). We found that explicit generalization was dominated by the extrinsic reference frame (70% extrinsic and 30% intrinsic; Figure 5A), while implicit generalization appeared to display an even mixture of extrinsic and intrinsic representations (51% extrinsic and 49% intrinsic; Figure 5B).

**Figure 5.**
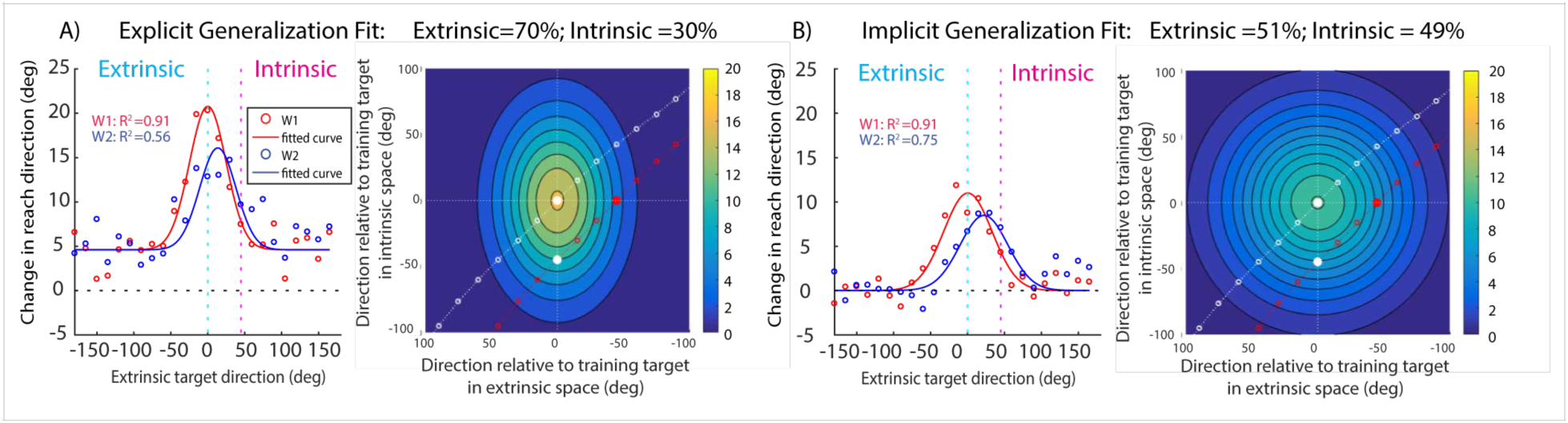
Extrinsic and intrinsic contributions to the explicit and implicit generalization patterns. Different mixture ratios for A) implicit and B) explicit forms of learning estimated by a gain-field model fit with varying extrinsic and intrinsic contributions to the data averaged across all subjects in W1 (red line) and W2 (blue line). Right panel: Representation of the gain-field model arising from the combination of extrinsic and intrinsic representations.

The bootstrap analysis confirmed the robustness of the preceding results. For explicit learning, the median contributions of the extrinsic and intrinsic reference frame were 70% (95% CI = 57.5%, 90%) and 30% (95% CI = 10%, 42.5%) respectively. In contrast, for implicit generalization the median extrinsic and intrinsic contributions were 50% (95% CI = 39%, 52%) and 50% (95% CI = 48%, 62%) respectively. These results show that each form of learning features distinct contributions from both intrinsic and extrinsic reference frames. Specifically, explicit learning was predominantly represented in an extrinsic reference frame, while implicit learning had a different ratio that had a more even mixture of extrinsic and intrinsic reference frames.

### Combination of explicit and implicit generalization pattern can account for the generalization pattern in the combined condition

With the generalization functions of explicit and implicit learning characterized in isolation, we then sought to determine if we could predict the generalization function when explicit and implicit learning combine in a standard visuomotor rotation task. To this end, the combined condition was designed such that subjects learned to compensate for the visuomotor rotation using a combination of explicit and implicit learning. To accomplish this, subjects were trained to compensate for a visuomotor rotation using online visual feedback, akin to a standard visuomotor rotation task. Aside from the difference in the form of visual feedback, the workspace locations and the locations of testing targets were identical to the explicit and implicit condition.

Subjects learned to compensate for the visuomotor rotation very well, displaying 26.9±1.1° of change in hand direction when averaged over the last 10 trials (Figure 6A). The asymptotic performance was significantly different from zero (t(15)=23.1, *p<0.0001*) and the amount rotation required to fully compensate for the visuomotor rotation (t(15)=-4.5, *p<0.0001*).

**Figure 6.**
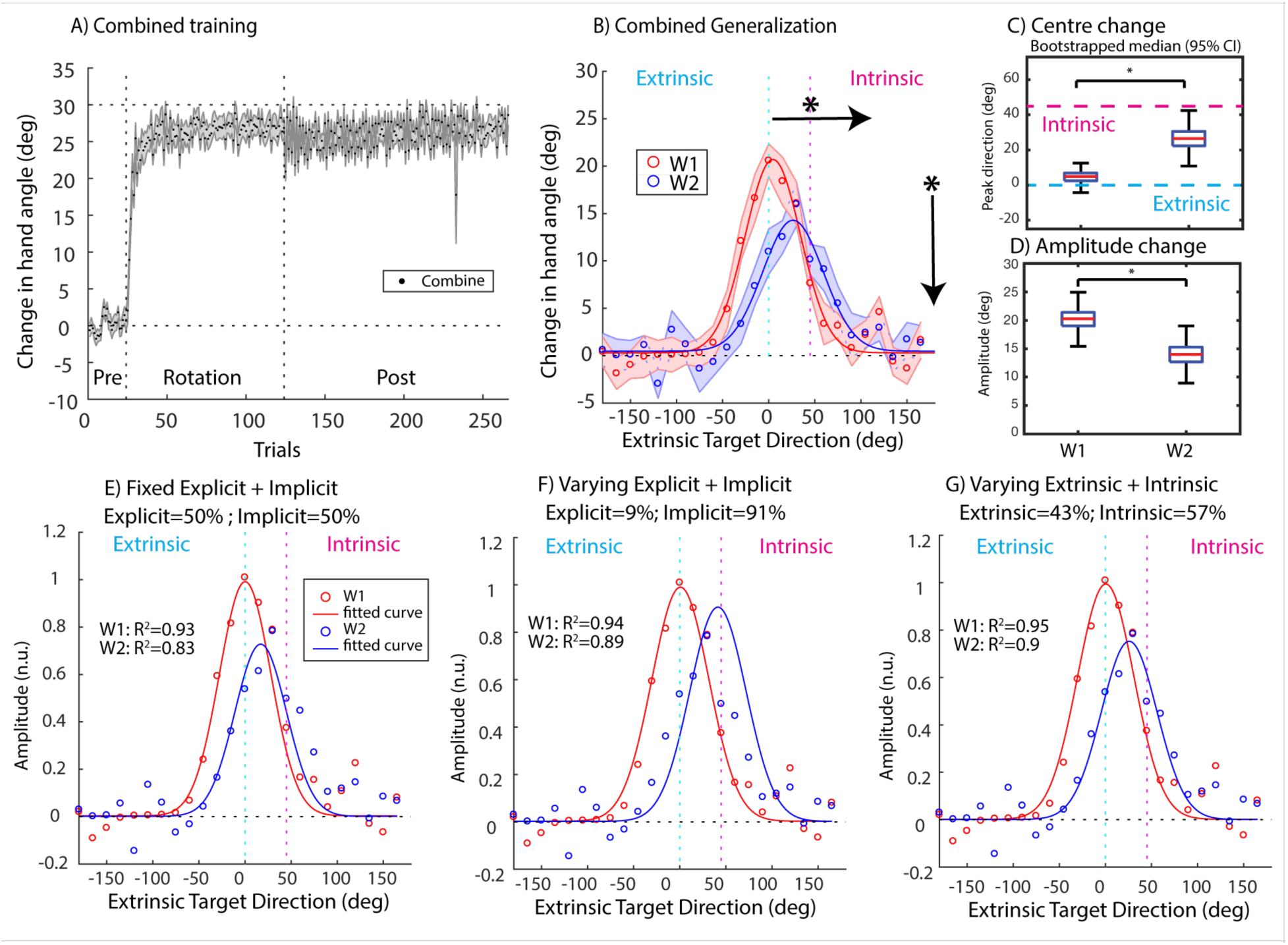
Gaussian fit to the generalization pattern at W1 and W2 in the combined condition (A) Mean change in hand angle when participant trained with a cursor that was aligned (Pre-rotation phase) or rotated 30° relative to hand movement (Rotation and Post-rotation phase) for the combined condition. B) Gaussian fits for the generalization pattern in the combined condition. The observed generalization pattern was averaged across all participants in each condition at W1 (red line) and W2 (blue line). Shaded area represents the mean±SE. C) Estimated center of generalization for each of the 2000 boot strap samples at W1 and W2 for the combined condition. D) Distribution of the amplitude of generalization for each of the 2000 boot strap samples at W1 and W2 for the combined condition. Quartiles and confidence intervals on indicated on the box plots. *Significant main effect following a bootstrap test at P <0.05. E) The generalization in W2 is best accounted for by an equal weighted sum of implicit and explicit components according to the BIC. F) Best fit to the subject averaged data when explicit and implicit weights allowed to vary. G) Best fit to the subject averaged data derived from a single process model that arises from a combination of extrinsic and intrinsic representations

The generalization function in the trained posture (W1) displays the typical Gaussian-like shape (Figure 6B). Indeed, the generalization pattern at W1 was characterized by a single gaussian with an amplitude of 22.8±1.6°, a center location of 5.3±7.3° with respect to the 0° training target, a width of 29.1±4.1° and an offset of 1.1±0.42°. When limb posture was rotated 45° to the transfer limb posture (W2), the generalization pattern maintained its characteristic Gaussian-like shape but with an apparent drop in the peak of generalization. We found that the generalization pattern had a peak that was centered at 25.1±7.8°, with an amplitude of 16.4±1.6°, a width of 35.2±3.2° and offset of 0.5±0.2. Comparison of the generalization functions at W1 and W2 revealed that the amplitude of generalization was significantly reduced (-6.4±2.8; t(15) = 3.5, p=0.003) and had a shift of 19.8±4.4° toward a pure intrinsic representation in W2. This phase shift was significantly different from the zero phase shift predicted by an pure extrinsic representation (t(15) = 4.3, p<0.0001) and the 45° phase shift predicted by the pure intrinsic representation (t(15) = -5.5, p<0.0001). In addition, there were no significant differences in the width (t(15) = 4.1, p=0.68) and vertical offset (t(15) = 1.4, p=0.17) of the generalization function between W1 and W2.

Interestingly, the reduction in amplitude and the degree of phase shifts are similar to that observed in the explicit and implicit conditions, respectively. This raises the possibility that the generalization pattern found in the combined condition could be attributed to a combination of generalization pattern arising from explicit and implicit learning. To test this idea, we fit a simple linear model whereby the overall generalization function is the result of a linear combination of the individual explicit and implicit generalization functions obtained in the explicit and implicit conditions, to the data in the combined condition. Remarkably, we found that, this model with no free parameters accounted for 83.4% of the variance in the generalization pattern at W2 for the combined condition (Figure 6E). Alternative models in which the weighting of the explicit and implicit components were allowed to vary (BIC = 0.6; Figure 6F) or single process models that arise from a combination of extrinsic and intrinsic representations (BIC =3.4; Figure 6G) did not produce a significantly better fit. These results suggest that the generalization pattern in a typical visuomotor rotation task arises from a combination of explicit and implicit forms of learning with distinct extrinsic and intrinsic contributions.

## Discussion

The goal of our study was to identify the reference frame used for representing explicit and implicit forms of learning, and to determine if their simple combination could explain previous reports of mixed representations in standard visuomotor rotation tasks. First, we observed that, when generalization was tested in W2, explicit learning was expressed predominantly in an extrinsic reference frame, but the amplitude of generalization was significantly reduced. In contrast, the implicit generalization function appeared to be phase shifted toward an intermediate position between the training target defined according to intrinsic and extrinsic reference frames when shifting to W2. Furthermore, implicit generalization did not reduce in amplitude when changing postures. These findings suggest that neither explicit nor implicit learning conform to a simple interpretation in terms of encoding in either a pure extrinsic or pure intrinsic reference frame. Thus, to gain further insights into this mixed representation, we fit a gain-field model which allows for varying contributions of extrinsic and intrinsic representations to explain the generalization function. We found that explicit generalization has a mixture ratio that is dominated by the extrinsic reference frame, while implicit generalization appeared to display an even mixture of intrinsic and extrinsic representations. Finally, we showed that a simple composition model whereby the overall generalization function is the result of a linear combination of explicit and implicit learning, is able to account for nearly 85% of the variance associated with the generalization pattern due to learning in a typical visuomotor task, when both processes are operating. Taken together, these findings suggest that each form of learning makes distinct contributions from both extrinsic and intrinsic reference frames, and the combination of these distinct features shapes the generalization pattern observed at novel limb postures.

### Explicit and implicit generalization in mixed reference frames

A number of studies have examined how learning in visuomotor rotation tasks generalizes across different limb postures and have found conflicting evidence. For example, Wang and Sainburg (2005) and Krakauer et al. (2000) have shown that learned compensations to visuomotor rotations are transferred in an extrinsic reference frame, while others have found evidence that learning may be represented in an intrinsic reference frame (Rotella et al. 2015; Krakauer et al, 2006; De Rugy et al 2009). There is also evidence that learning takes place in both extrinsic and intrinsic reference frames (Brayanov et al. 2012; Poh et al. 2017). Brayanov et al. (2012) probed generalization across a wide distribution of movement directions at various limb postures and showed that learning generalized most strongly to a target direction that is intermediate between the intrinsically and extrinsically defined training direction. This pattern of generalization was found to be most consistent with a multiplicative gain-field model with a mixture of extrinsic and intrinsic representations. We hypothesized that the failure to find a consistent representation is the result of the concurrent operation of explicit and implicit learning processes, each operating within a distinct representational space. As a consequence, different patterns of generalization in extrinsic, intrinsic or mixed reference frames will arise depending on the relative contribution of generalization effects driven by independent effects from both explicit and implicit forms of learning (Heuer and Hegele, 2008, 2011; McDougle et al. 2017).

To assess this possibility, we measured the individual generalization functions associated with explicit and implicit learning in relative isolation using a modified visuomotor rotation task. To this end, we hypothesized that explicit forms of learning may reflect a deliberate strategy to re-aim movements toward the training target, and thus be related to an extrinsic reference frame. In contrast, implicit forms of learning may depend more on an intrinsic reference frame since adaptive changes to motor output may be more related to limb posture and joint configurations. However, our behavioral results did not fit this view. Specifically, while explicit learning was expressed predominantly in an extrinsic reference frame, the amplitude of generalization decayed with modulations in limb posture, which complicates a pure extrinsic reference frame interpretation. In comparison, while the amplitude of implicit generalization did not decay substantially with changes in limb posture, the direction of peak generalization was phase shifted to an intermediate position between the extrinsically and intrinsically defined training direction, reflecting a mixed representation. These findings suggest that the representation underlying explicit and implicit learning may not rely on an easily categorizable extrinsic or intrinsic reference frame as we had initially theorized.

### Implications for explicit learning

For explicit learning, we showed that changing limb posture did not strongly influence the direction in which learning was expressed; instead the amplitude of generalization was reduced. This suggests the amplitude of generalization is influenced by intrinsic limb position based learning effects. One possible interpretation could be that limb posture might serve as a contextual cue that is coupled with the explicit learning such that generalization is reduced when limb posture is altered (Baraduc and Wolpert 2002; Gandolfo et al. 1996; Howard et al. 2012; Howard et al. 2013; Krakauer et al. 2006). The idea that the extent of generalization of learning between movements depends on the similarity of the underlying intrinsic joint-based requirements is reflected in previous studies that have shown that extent of generalization is tied to specific muscles (de Rugy et al. 2009; Poh et al. 2017; Krakauer et al. 2006), and joint postures (Baraduc and Wolpert 2002) involved during training. Our findings are consistent with this work by showing that the intrinsic limb-posture dependent reduction in the amplitude of generalization is limited to explicit learning, but not implicit learning.

It is important to note that our generalization task was designed to isolate explicit and implicit learning using different visual feedback manipulations, which allowed us to independently examine how each form of learning generalize to movements in the untrained limb posture. Our approach differs from other studies where explicit estimations of the movement direction required to compensate for the visuomotor rotation were obtained to quantify the relative contributions of explicit and implicit forms of learning. For example, in the study by Heuer and Hegele (2011), following learning of the rotated visual feedback, subjects verbally instructed the experimenter to rotate a guide line to align with their intended movement direction to compensate for the perturbation. While others studies have instructed participants to verbally state their aiming direction with reference to a numbered landmark during learning (Bond and Taylor 2015; Taylor et al. 2014, McDougle et al. 2017). Such re-aiming strategies have been interpreted as an explicit form of learning that the motor system implements in parallel with implicit learning, and the difference between the direction of aim and the actual reach direction constitutes a measure of implicit learning. However, studies employing these approaches have often noted that explicit learning results in broad generalization across all movement directions (Heuer and Hegele 2008, 2011; McDougle et al. 2017). As such, it is challenging to determine which reference frames contribute to the representation of explicit learning because the learning is not direction-specific and the generalization effects in extrinsic or intrinsic reference frames are indistinguishable at the untrained posture.

Unlike in those studies, here we have found relatively narrow generalization of explicit learning around the training direction in W1. When generalization was tested to W2, the pattern of explicit generalization was most consistent with an intermediate representation that is predominantly expressed in the extrinsic reference frame, and that the amplitude of generalization is modulated by changes in limb posture. The difference between our findings and prior approaches is intriguing, and at present, its unclear. One possible explanation is that explicit estimations of the aiming direction may encourage subjects to “solve” for the rotation by developing a strategy to re-aim their movements in a direction opposite to the rotation for every target. Such a phenomenon may bear resemblance to the “Hawthorne Effect”, commonly observed in social psychology (Franke 1980; Franke and Kaul 1978). That is, by simply asking participants what they are doing, they necessarily change the way they respond. Thus, it is unclear if the broad generalization function of explicit aiming observed in previous studies reflects the true mark of explicit learning or an artifact of task instructions.

### Implications for implicit learning

Our findings show that extent of implicit generalization was not modulated by the change in limb posture, but the peak direction of generalization was shifted to an intermediate direction between the trained target location defined in an intrinsic and an extrinsic reference frame. This shift in the generalization pattern resembles the shift in the generalization pattern that was reported by Brayanov et al. (2012), and that has been taken as evidence that learning is represented in a multiplicative gain-field combination of intrinsic and extrinsic movement representations. However, because the authors did not dissociate between explicit and implicit learning in their study, it was difficult to attribute the shift in the generalization pattern in the proposed mixed reference frame to either the explicit or the implicit form of learning. Given that we have designed our experiments to isolate effects of explicit and implicit forms of learning across different limb postures, the observed shifts in implicit generalization only suggest that the shift in the generalization in Brayanov et al (2012) was primarily due to an implicit form of learning. By this view, implicit learning could involve an integrated representation which combines both extrinsic and intrinsic reference frames. Taken together, the involvement of multiple reference frames in both explicit and implicit learning is consistent with previous modeling studies, which suggest that learning might occur concomitantly in multiple reference frames. However, the amplitude and shape of generalization expressed in a particular reference frame is determined by how the motor system attribute error to sensory signals in different reference frames (Berniker and Kording 2008; Berniker and Kording 2011).

### Conclusion

In conclusion, our results demonstrate that generalization of explicit and implicit learning did not conform to a simple interpretation in terms of encoding in a categorical extrinsic or intrinsic reference frame. While explicit learning was expressed predominantly in an extrinsic reference frame and decays with changes in limb posture, implicit generalization appeared to be phase shifted toward an intermediate position between the training target according to intrinsic and extrinsic reference frames and does not decay substantially with postural changes. Further, we showed that a simple linear model whereby the overall generalization function is the result of a linear combination of explicit and implicit generalization observed captures nearly 85% of the variance in the generalization pattern in a standard visuomotor rotation task, when the two processes are operating. These results suggest that multiple reference frames contribute to each form of learning, and the combination of these distinct features shapes the generalization pattern observed at novel limb postures. While our results suggest that the representation underlying explicit and implicit learning may not rely on an easily categorizable extrinsic or intrinsic reference frame, it remains an open question as to why both explicit and implicit learning arise from mixed representations. One intriguing possibility is that learning is not truly represented in an extrinsic or intrinsic reference frame, but rather is learned in a higher dimensional representation and we are simply observing low-dimensional projections from this higher representation (Shepard 1987).

## Acknowledgements

We would like to thank Carlo Campagnoli for helpful discussions and comments on the manuscript. This work was supported by the National Institute of Neurological Disorders and Stroke (Grant R01 NS-084948) and the National Science Foundation under Grant No. 1838462.

